# Advanced understanding of prokaryotic biofilm formation using a cost-effective and versatile multi-panel adhesion (mPAD) mount

**DOI:** 10.1101/2021.09.09.459712

**Authors:** Stefan Schulze, Heather Schiller, Jordan Solomonic, Orkan Telhan, Kyle Costa, Mechthild Pohlschroder

## Abstract

Most microorganisms exist in biofilms, which comprise aggregates of cells surrounded by an extracellular matrix that provides protection from external stresses. Based on the conditions under which they form, biofilm structures vary in significant ways. For instance, biofilms that develop when microbes are incubated under static conditions differ from those formed when microbes encounter the shear forces of a flowing liquid. Moreover, biofilms develop dynamically over time. Here, we describe a cost-effective, 3D-printed coverslip holder that facilitates surface adhesion assays under a broad range of standing and shaking culture conditions. This multi-panel adhesion (mPAD) mount further allows cultures to be sampled at multiple time points, ensuring consistency and comparability between samples and enabling analyses of the dynamics of biofilm formation. As a proof of principle, using the mPAD mount for shaking, oxic cultures, we confirm previous flow chamber experiments showing that *Pseudomonas aeruginosa* wild type and a phenazine deletion mutant (Δ*phz*) form similar biofilms. Extending this analysis to anoxic conditions, we reveal that microcolony and biofilm formation can only be observed under shaking conditions and are decreased in the Δ*phz* mutant compared to wild-type cultures, indicating that phenazines are crucial for the formation of biofilms if oxygen as an electron acceptor is not available. Furthermore, while the model archaeon *Haloferax volcanii* does not require archaella for attachment to surfaces under static conditions, we demonstrate that *H. volcanii* mutants that lack archaella are negatively affected in their early stages of biofilm formation under shaking conditions.

**Importance:** Due to the versatility of the mPAD mount, we anticipate that it will aid the analysis of biofilm formation in a broad range of bacteria and archaea. Thereby, it contributes to answering critical biological questions about the regulatory and structural components of biofilm formation and understanding this process in a wide array of environmental, biotechnological, and medical contexts.

## Introduction

Planktonic prokaryotic cells often encounter highly stressful environmental conditions such as those created by toxins or nutrient depletion. One strategy that bacteria and archaea have evolved to mitigate such stress is the establishment of a biofilm: a complex microbial community bound by a matrix of extracellular polymeric substances. The first steps in biofilm formation are cell adherence to an abiotic surface quickly followed by cell aggregation and microcolony formation; later, a mature biofilm is established. Adhesion often requires surface filaments (1–3). For example, *Pseudomonas aeruginosa*, a bacterium in which biofilm formation has been thoroughly studied, requires flagella and type IV pili to complete the initial steps of establishing a biofilm (4). For some organisms, additional components are also involved in biofilm maturation, such as redox-active secondary metabolites called phenazines. In *P. aeruginosa*, phenazines drive the release of extracellular DNA into the biofilm matrix and facilitate survival of cells in the anoxic core of a biofilm (5, 6).

The large variety of platforms that have been developed to study biofilm formation seemingly reflects the diversity of attachment mechanisms and biofilm architectures that have evolved. While evolutionarily conserved type IV pili are required for surface adhesion in many organisms (7), as analyzed e.g. through simple atmosphere-liquid interface (ALI) assays in standing cultures, the use of specific pilins can vary depending on the environmental conditions (2, 8, 9). Furthermore, while bacterial flagella and their archaeal counterparts (archaella) facilitate the binding to surfaces in some species (10), surface attachment in other species like the model haloarchaeon *Haloferax volcanii* appears to be independent of archaella (11). However, the involvement of type IV pili, archaella, and other cell surface structures in attachment and biofilm formation may depend on the environmental conditions under which ALI assays are performed. In fact, it has been shown that shear forces impact the ability of microbes to form biofilms, the characteristics of the established biofilms, and the rate of detachment in prokaryotic biofilms, e.g. for *Escherichia coli* (12). Therefore, it is critical to analyze the formation of biofilms under various conditions, and more advanced platforms, such as complex flow chambers or rotating annular bioreactors, have been used for this task (13).

Despite the importance of generating data that will lead to critical discoveries about the molecular mechanisms that regulate the establishment of biofilms, each biofilm analysis method has intrinsic limitations. For instance, the standard ALI assay cannot be used to evaluate biofilm formation under flow (14) and thus prevents insight into the effect of shear forces on biofilm formation. Furthermore, this type of standing assay often allows undesirable liquid biofilm formation (15–17), making comparisons of planktonic and sessile cells unfeasible. As an alternative, some simple assays like the BioFilm Ring Test may be performed under shaking conditions, but this advantage comes with the drawback that microscopy cannot be used to analyze the biofilm (14). In contrast, more complex systems like flow chambers allow for detailed continuous imaging of biofilms (13, 18); however, it is extremely difficult to retrieve samples for molecular biological or biochemical evaluation at various time points. Some of these systems, including flow chambers and rotating annular bioreactors, are also costly (13). These limitations highlight that most biofilm analysis methods face a tradeoff between high throughput and detail of analysis. High-throughput platforms like the standard ALI assay or BioFilm Ring Test only allow for rather superficial analyses of biofilm formation, while detailed analyses, e.g. in flow chambers, suffer from comparatively low throughput and limited versatility.

To overcome some of these challenges, we present a detailed description of the use of an inexpensive **m**ulti-**p**anel **ad**hesion (mPAD) mount for the characterization of biofilm formation. This mount, which can be produced using a standard 3D printer, facilitates the analysis of cell adhesion to coverslips for both standing and shaking cultures and under a variety of conditions. Using the mPAD mount, the same culture can be sampled at multiple time points for microscopic, biochemical, and molecular biological analyses.

As a proof of principle, we have analyzed *P. aeruginosa* biofilm formation under standing and shaking conditions, showing not only differences in biofilm architecture and the ability to proceed past the single-cell adhesion stage under anoxic conditions but also a decrease in microcolony formation in a Δ*phz* mutant strain under shaking, anoxic conditions in contrast to wild-type cultures. Similarly, *H. volcanii* biofilms that formed in shaking cultures look very different from biofilms established in standing cultures, and we show that the Δ*arlA1/2* strain exhibits a decreased initial surface adhesion in shaking cultures despite lacking any defects under standing conditions, highlighting the importance of testing biofilm formation in wild-type and mutant strains under a variety of conditions.

## Materials and Methods

### Strains and growth conditions

*P. aeruginosa* PA14 and the *P. aeruginosa* Δ*phz* mutant (19) were grown at 37 °C in lysogeny broth (orbital shaker at 250 rpm). For anaerobic growth, 40 mM sodium nitrate and 200 mM MOPS were included in the medium, and cultures were grown standing inside of a heated (37 °C) Coy-type anoxic chamber with an atmosphere of 2-3% H_2_, 10% CO_2_, and balance N_2_. MOPS buffer was included to avoid pH changes from reduction of nitrate and the high CO_2_ atmosphere in the chamber. For growth as a biofilm, overnight cultures were sub-inoculated, grown for several hours to an OD_600nm_ between 1.0 and 1.5, and then further diluted to an OD_600_ of 0.3. These cultures were immediately transferred to biofilm growth conditions and incubated either standing in 12-well plates or shaking in Petri dishes (100 mm x 15 mm) with the mPAD mount (see below).

*H. volcanii* H53 as well as the *H. volcanii* Δ*arlA1/2* mutant were grown at 45 °C in liquid (orbital shaker at 250 rpm) or on solid agar (containing 1.5% (wt/vol) agar) semi-defined casamino acid (Hv-Cab) medium (20), supplemented with tryptophan (+Trp) and uracil (+Ura) at a final concentration of 50 μg ml^−1^. A colony from a solid Hv-Cab plate (incubated for three to five days at 45 °C) was inoculated into 5 mL Hv-Cab liquid medium. After incubating the culture tubes overnight at 45 °C with shaking (orbital shaker at 250 rpm) until the strains reached mid-log phase (OD_600nm_ ~0.7) the strains were diluted to an OD_600nm_ of 0.01 into a final volume of 20 mL followed by incubation at 45 °C with shaking (orbital shaker at 250 rpm) until cultures reached an OD_600nm_ of 0.35. Each culture was then transferred into either a 12-well plate or a sterile plastic Petri dish (100 mm x 15 mm) for biofilm growth conditions (see below).

### Printing of mPAD mount

The 3D model of the mPAD mount can be downloaded from Thingiverse (https://www.thingiverse.com/thing:4784964). A Form 2 (Formlabs) 3D printer using a High Temp resin (Formlabs) was used to print mPAD mounts that can be sterilized in an autoclave. These mounts were used for the *P. aeruginosa* biofilm assays. Conversely, an MP Mini Delta 3D printer (Monoprice) using polylactic acid (PLA) filament (Monoprice) was employed to 3D-print the mPAD mounts used for assays with *H. volcanii*.

### Setting up the mPAD mount for adhesion assays under standing and shaking conditions

*P. aeruginosa* biofilms: glass coverslips (22 mm x 22 mm x 1.5 mm, Fisher Scientific) were marked to be able to distinguish the two sides (facing inward and outward from the mPAD) as well as the top and bottom of the coverslip. When inserting the coverslips into the mPAD mount, thin pieces of aluminum foil were placed at the top of the coverslip to enable a tighter fit. Assembled mPADs were autoclaved to sterilize all surfaces. *P. aeruginosa* culture (grown and diluted as described above) was placed in the bottom half of a Petri dish, and the mPAD mount was placed on top of the Petri dish with each coverslip reaching at least 0.5 cm into the culture. The Petri dish lid was placed on top of the mPAD mount before incubating the assembled setup in a humidified chamber at 37 °C. Cell attachment was allowed to occur for 30 minutes before placing the chamber on an orbital shaker set to 60 rpm. For static biofilms, 2 ml of culture at OD_600nm_ 0.3 was aliquoted into a 12-well plate (Falcon) with a glass coverslip placed into each well. Both wild-type and Δ*phz* biofilms were grown via the methods described above under both oxic and anoxic conditions.

*H. volcanii* biofilms: mPAD mounts were washed with 70% ethanol and allowed to air dry before inserting the marked coverslips (see above) into the slits of the mPAD mount. Thin pieces of autoclaved aluminum foil were placed at the top of the coverslip to enable a tighter fit in the mPAD mount. The mPAD mount with inserted coverslips was placed onto the bottom half of the Petri dish containing the liquid culture with each coverslip reaching at least 0.5 cm into the culture. The Petri dish lid was placed on top of the mPAD mount before placing the Petri dish with the mPAD mount into a humidified chamber. The cultures were incubated at 45 °C on an orbital shaker set to 60 rpm. For static biofilms, 2 ml of *H. volcanii* culture at OD_600nm_ 0.35 was aliquoted into a 12-well plate (Falcon) with a glass coverslip placed into each well. Both wild-type and Δ*arlA1/2* biofilms were grown via the methods described above under oxic conditions.

### Staining and imaging

*P. aeruginosa* biofilms: coverslips were removed at 1, 6, 20, and 45 hours, submerged briefly in 0.5% NaCl to wash off loosely attached cells, and fixed by placing the coverslip in 3 ml of a 0.5% NaCl/2% glutaraldehyde solution in a 12-well plate for 30 minutes at room temperature. For anaerobically grown cultures, the fixation step was performed inside the anoxic chamber before moving coverslips into an oxic atmosphere for further processing. After fixation, the coverslips were immediately placed into 3 ml of 0.1% crystal violet for 10 minutes. After staining, coverslips were washed in H2O and allowed to dry before imaging. Coverslips were imaged on an ECHO Revolve R4 hybrid microscope with a universal slide mount. The microscope was operated in the upright orientation with an extra-long working distance condenser (NA 0.30, working distance 73 mm). Images were taken with either a 20x fluorite (NA 0.45, working distance 6.6-7.8 mm) or 40x fluorite (NA 0.75, working distance 0.51 mm) objective lens. Images were collected from the middle of the slide near the location of the ALI.

*H. volcanii* biofilms: coverslips were removed at 24 and 120 hours, submerged briefly in 18% saltwater, and fixed by placing them in 2% acetic acid in small Petri dishes (Corning) for 30 minutes at room temperature. After fixation, the coverslips were allowed to air dry. Cells attached to coverslips were stained by submerging them into 0.1% crystal violet for 10 minutes. After removing the coverslips from the crystal violet solution, they were rinsed twice with H_2_O and air dried. Biofilms were imaged using a Leica DMi8 inverted microscope with brightfield settings using 20x and 40x magnifications. Representative images of all areas of the biofilm on the coverslip were taken. Images were collected from the middle of the slide near the location of the ALI.

## Results and Discussion

### The mPAD mount is designed as a versatile and affordable device to characterize biofilm formation

Many invaluable tools have been developed over the past decades to characterize biofilm formation, ranging from simple ALI assays on coverslips in 12-well plates to sophisticated flow chambers that allow live observation of the stages of biofilm formation and comparison of various shear forces. To combine advantages of different types of assays, we designed the mPAD mount (Fig. 1). 3D-printing of the mPAD mount results in low-costs, similar to simple 12-well plate assays, and its design for standard Petri dishes ensures its adaptability to different environmental conditions, including shear forces generated by constant shaking on an orbital shaker and an O_2_-deprived atmosphere in an anaerobic chamber. Additionally, multiple slots within each mPAD mount enable the observation of different stages of biofilm formation by removing individual coverslips at different time points. This design also allows harvesting of samples for transcriptomics or proteomics analyses from different stages of biofilm formation from a single culture, thereby reducing high variabilities that can arise from the use of multiple cultures. Furthermore, when used under shaking conditions, the formation of liquid biofilms, which can obstruct comparative system-wide analyses because of varying planktonic subpopulations, is prevented.

**Figure 1:**
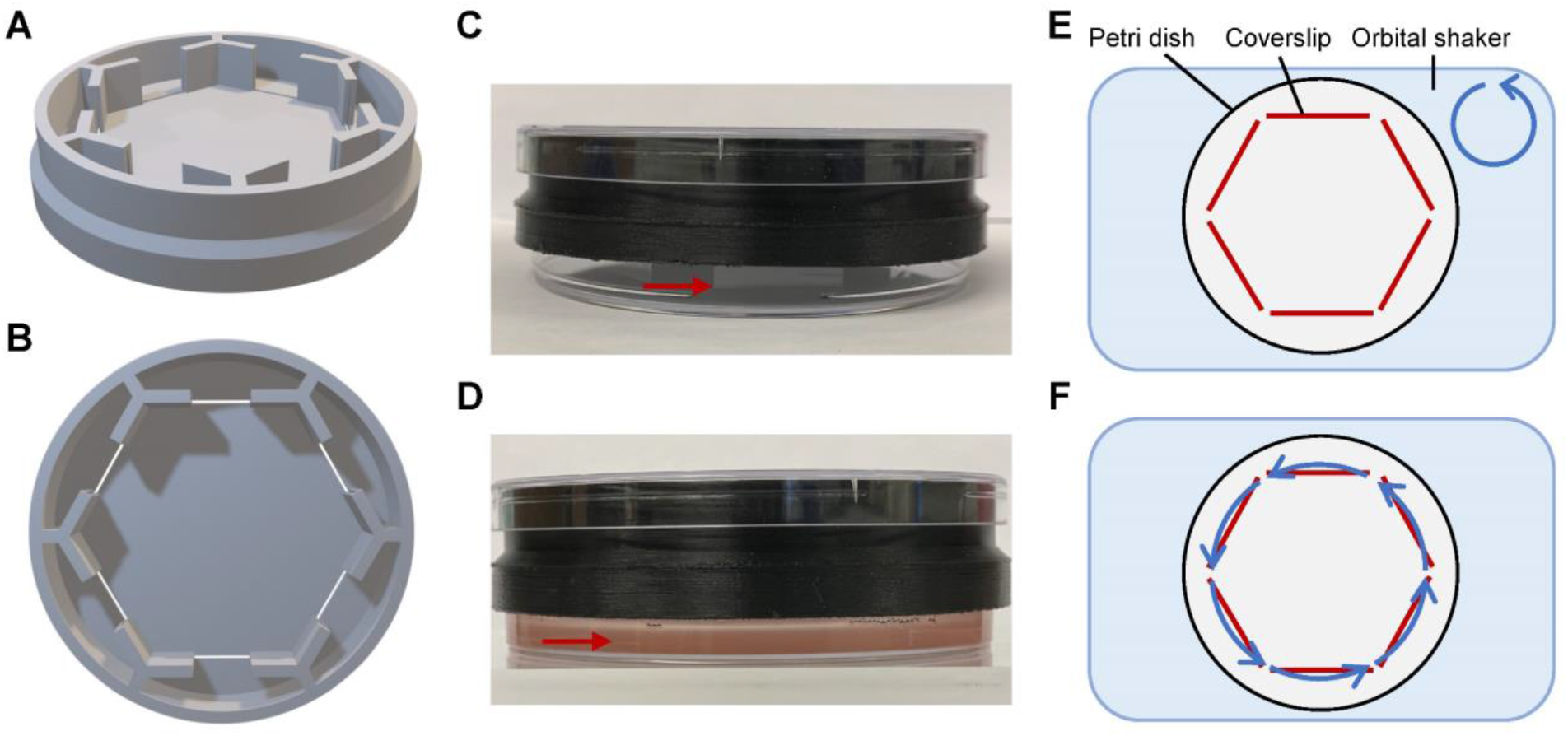
The 3D-printed mPAD mount can be used for biofilm assays under a broad range of conditions, including shaking. A 3D-model of the mPAD mount is shown from the side (**A**) and top (**B**). Coverslips can be inserted into the six slots of the mPAD mount and subsequently placed onto a Petri dish (**C**). After filling the Petri dish with a cell culture (*H. volcanii* shown here), coverslips reach into the culture (**D**), allowing cells to adhere to the coverslip surface. The mPAD mount can be placed on an orbital shaker to expose cells to shear forces (**E**). The radial positioning of the slots of the mPAD mount ensures that cells adhering to any coverslip experience the same shear forces (**F**). Red arrows point to the lower end of inserted coverslips; blue arrows indicate shaking of the orbital shaker and the shear forces experienced by the coverslips.

Radial positioning of the six slots of the mPAD mount ensures that all coverslips are exposed to the same shear force when placed on an orbital shaker (Fig. 1E-F). With this design only minimal differences between the inner and outer side of the coverslip were observed (data not shown). However, due to the 3D-printing of the mount, changes to the design are straight-forward, and a variety of different layouts are conceivable, e.g. to increase the adhesion surface area or to accommodate movement of the liquid via a stirring bar instead of an orbital shaker (Supplemental Figure 1).

3D-printing of the mPAD furthermore supports a wide variety of materials, including materials that show high resistance to chemicals and temperature. In this study, we have used a Form 2 (Formlabs) 3D printer employing a High Temp resin (Formlabs) for the *P. aeruginosa* studies, as that material allows the mPAD mounts to be sterilized in an autoclave. For the biofilm studies of the archaeon *H. volcanii*, an MP Mini Delta 3D printer (Monoprice) using polylactic acid (PLA) filament (Monoprice) was employed to 3D-print the mPAD mount. While PLA is not as resistant to heat and chemicals as Formlabs’ High Temp resin, it is more affordable, and contamination of *H. volcanii* mid-log phase cultures grown in media containing ~2.5 M salt are very unlikely, thus semi-sterile 70% ethanol washes could be employed instead of autoclaving.

### The mPAD mount is suitable for characterization of *P. aeruginosa* wild-type and *Δphz* biofilms exposed to shear forces

The use of flow chambers has established that cell exposure to shear forces plays a significant role in the structure and architecture of *P. aeruginosa* biofilms (21). Similarly, employing the mPAD mount setup on an orbital shaker facilitated the visualization of shear force effects on *P. aeruginosa* cultures, because cells adhering to coverslips within this setup are experiencing shear forces from the moving liquid. For wild-type *P. aeruginosa* cultures, adhesion of individual cells to the coverslip after an hour was similar under shaking conditions as under standing conditions using a common 12-well-plate ALI assay (Fig. 2). At six hours, microcolonies were observed, and by 20 hours, mature biofilms had formed under both conditions, but the biofilms formed under standing conditions were distributed more uniformly in a mixture of single cells, microcolonies, and large, three-dimensional biofilm structures. In contrast, under shaking conditions, cells formed large, dense, web-like structures with fewer visible unconnected microcolonies and single cells. The Δ*phz* strain showed a similar trend, although adhesion, microcolony formation, and mature biofilms were overall less dense compared to the wild type.

**Figure 2:**
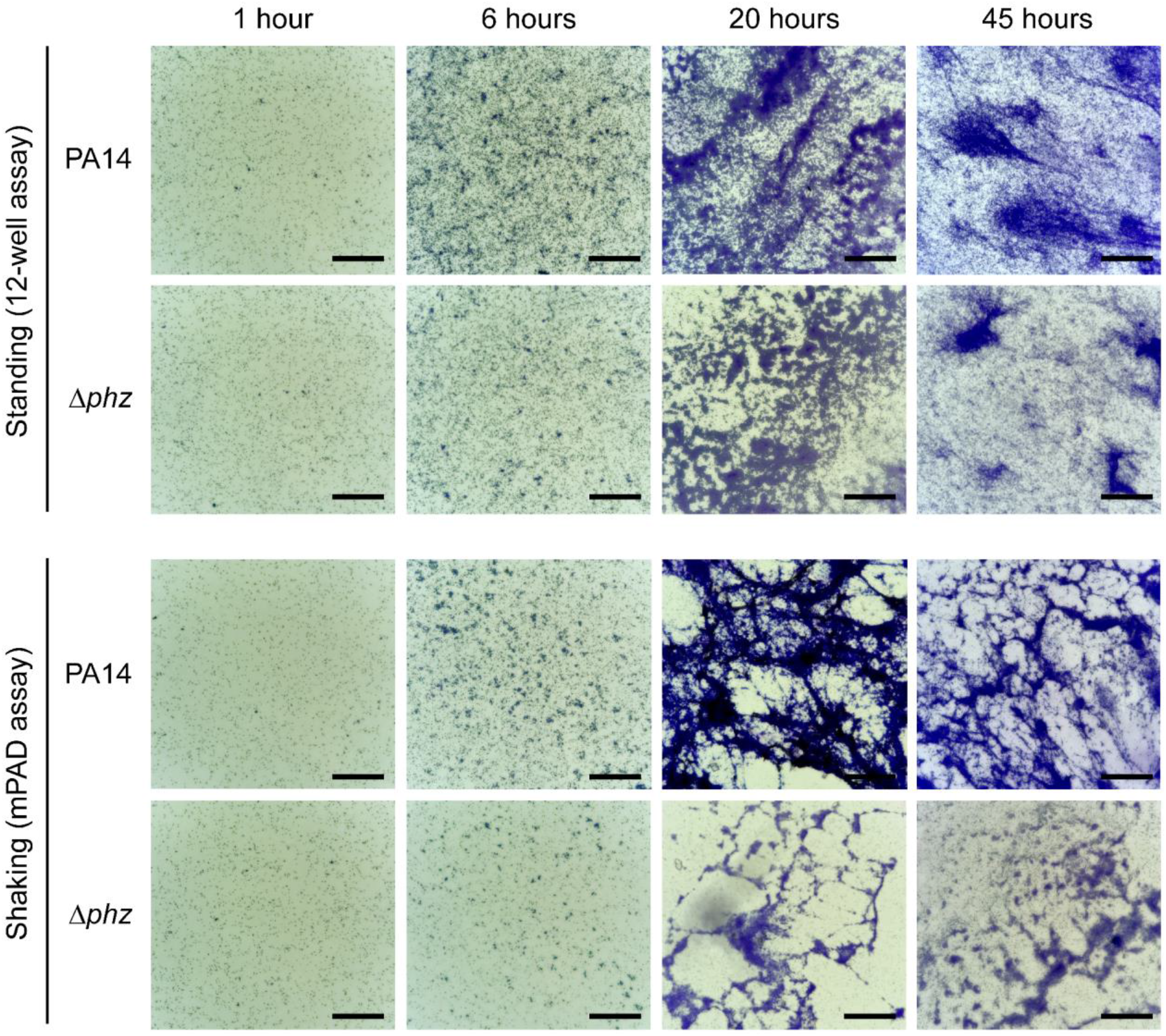
Using the mPAD mount for the analysis of *P. aeruginosa* biofilms under aerobic, shaking conditions confirmed that exposure to shear forces leads to different biofilm architectures and noticeable phenotype in Δ *phz* mutant. *P. aeruginosa* wild-type (PA14) and Δ*phz* cultures were incubated under aerobic standing and shaking conditions for 1, 6, 20, and 45 hours before staining adherent cells with crystal violet. Images are representative of two biological replicates. Scale bar is 100 ◻m.

The results show that the mPAD mount is suitable for the analysis of different stages of biofilm formation under shaking conditions, with differences in biofilm architecture that are in line with previous experiments using flow chambers (6). For further analyses, cells can be washed off the coverslips, e.g. to quantify adhesion through OD_600nm_ measurements of crystal violet. Untreated adhering cells can also be retrieved, e.g. for -omics experiments. Thereby, the mPAD mount sets the stage to compare the proteomes of biofilms formed under standing and shaking conditions to determine whether the differences are mainly structural or also include changes in the composition of the biofilm, including differentially expressed and post-translationally modified proteins or differences in the exopolymeric substances.

### Under anaerobic conditions, *P. aeruginosa* only forms microcolonies when exposed to shear forces

*P. aeruginosa* wild-type cultures can grow anaerobically, but comparisons of biofilm formation with and without the exposure to shear forces have thus far not been reported. In addition to characterizing wild-type biofilm formation under anaerobic, standing conditions, the versatility of the mPAD mount facilitated the analysis of shear force impacts on surface attachment, microcolony formation, and biofilm formation under anaerobic conditions. Interestingly, while initial attachment to the coverslip occurred both with and without shaking, only cells exposed to shear forces through orbital shaking in the mPAD assay were able to form microcolonies and biofilms without available O2, although these biofilms were significantly less dense than those under aerobic conditions (Fig. 3). Similar to the wild type, the Δ*phz* mutant was unable to form microcolonies under anaerobic, standing conditions. However, in comparison to the wild type, the adhesion and microcolony formation after 20 and 45 hours under anaerobic conditions were reduced for Δ*phz* and even lacking completely in one of the replicates. While this phenotype is similar to the reduced biofilm formation under aerobic conditions, the more pronounced effect under anaerobic conditions indicates a crucial role of *phz* as an electron acceptor when O2 is not available. This role is in line with results on electron shuttling via phenazines in *P. aeruginosa* biofilms on agar plates (22).

**Figure 3:**
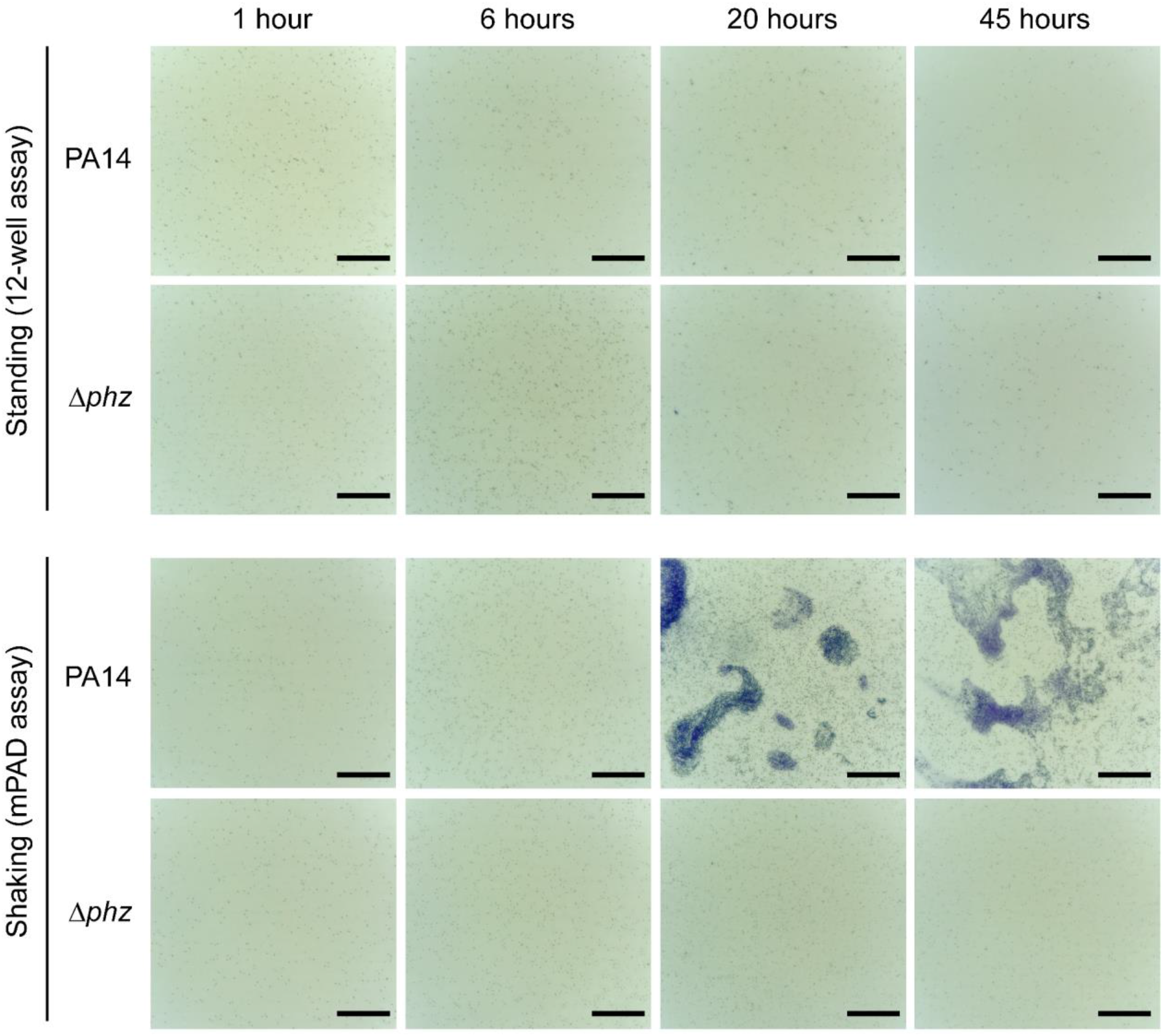
Employing the mPAD under anoxic conditions revealed that shear forces contribute to the formation of biofilms, which are dependent on *phz* under these conditions. *P. aeruginosa* wild-type (PA14) and Δ*phz* cultures were incubated in an anaerobic chamber under standing and shaking conditions for 1, 6, 20, and 45 hours before staining adherent cells with crystal violet. Images are representative of two biological replicates. Scale bar is 100 ◻m.

### *Haloferax volcanii* archaellins contribute to effective initial adhesion when exposed to shear forces

While the haloarchaeon *H. volcanii* has been used as a model organism to study biofilm formation in archaea (15, 16, 23), to our knowledge, biofilm assays have not been performed under shaking conditions for any archaeon so far. Using the mPAD assay under shaking conditions, we could show that, similar to *P. aeruginosa, H. volcanii* can form biofilms under shaking conditions but exhibits distinct biofilm architectures depending on the exposure to shear forces (Fig. 4). Under standing conditions, after 24 hours, attachment of *H. volcanii* to the coverslip surface resulted in a dense layer of cells interspersed with areas of small microcolonies, which developed into larger areas of dense biofilms within 120 hours. In contrast, under shaking conditions, *H. volcanii* attached only sparsely as single cells to the coverslip within 24 hours, and microcolonies formed as dense clusters of cells with network-like connections that likely consist of extracellular substances. After 120 hours, these microcolonies grew into dense biofilm structures that were connected over larger areas.

**Figure 4:**
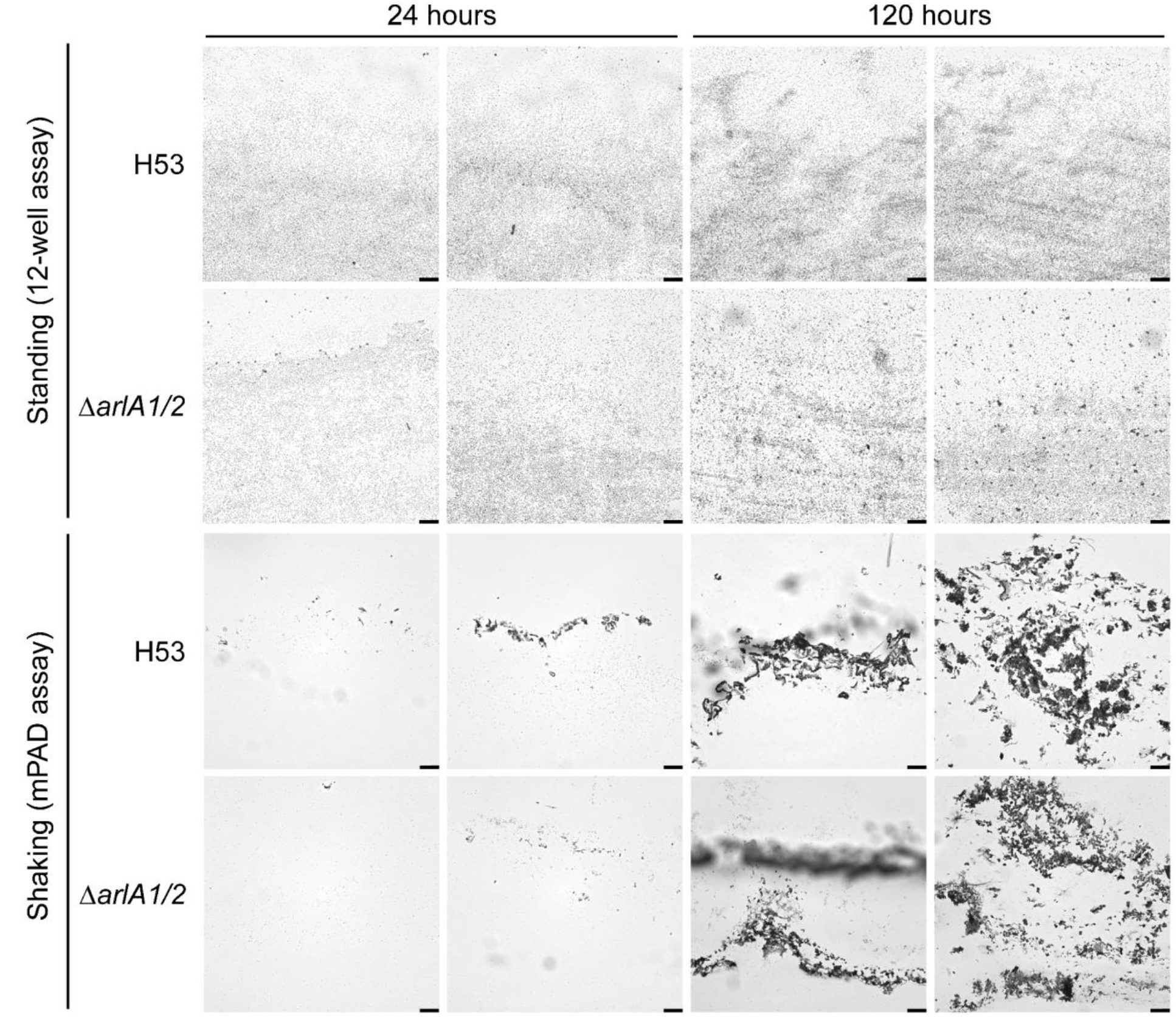
*H. volcanii* biofilm formation under shaking conditions differs substantially from standing conditions and is negatively affected by a lack of archaella. *H. volcanii* wild-type (H53) and Δ*arl1/2* cultures were incubated under standing and shaking conditions at 45 °C for 24 and 120 hours before staining adherent cells with crystal violet. Images are representative of two biological replicates. Scale bar is 50 ◻m.

The process of cell adhesion and microcolony formation without exposure to shear forces is independent of archaella in *H. volcanii*, as shown by the wild type-like adhesion of a Δ*arlA1/2* mutant, which lacks both archaellins (11). This observation is in contrast to many bacteria and archaea for which flagella and archaella, respectively, have been shown to be required for adhesion to surfaces (1, 24–27). To determine whether *H. volcanii* archaellins are required for biofilm formation when cells are exposed to shear forces, we compared the attachment of *H. volcanii* wild-type and Δ*arlA1/2* cultures to coverslips under shaking conditions using the mPAD mount. At 120 hours, both strains were able to form microcolonies and mature biofilm structures of similar magnitude and density (Fig. 4). However, at earlier stages, the mutant Δ*arlA1/2* strain did not adhere to the same extent as the wild type. While wild-type cultures at 24 hours already formed microcolonies in addition to single cell adhesion, Δ*arlA1/2* exhibited mainly single cell adhesion, and only few, small clusters of cells were present.

These results indicate that archaella in *H. volcanii* could be involved in the initial surface attachment under shaking conditions. Possible explanations for a contribution of archaella in the binding to surfaces could be an increased sensing of the surface or movement toward it.

In general, in contrast to bacteria, the effects of shear forces on archaeal surface attachment and biofilm formation have not been analyzed so far. Our initial results here suggest that valuable insights into the molecular mechanisms of this process in archaea could be gained. The mPAD mount provides an ideal foundation for further studies because it can be used in a broad range of extreme conditions in which many model archaea thrive.

## Conclusion

Biofilms represent the major life form of prokaryotes on Earth (28), and the conditions under which they occur are highly diverse and often not static. Therefore, we developed the simple, 3D-printed, versatile and affordable mPAD mount, which can be used to study prokaryotic biofilm formation under a broad variety of environmental conditions. The design of the mPAD thereby allows for samples to be taken from the same culture at multiple time points for microscopy as well as -omics analyses, capturing dynamic changes within the process of biofilm formation.

We could show the usefulness of the mPAD on the examples of *P. aeruginosa* and *H. volcanii* biofilm formation. For *P. aeruginosa*, we revealed that while biofilms do not form in the absence of O_2_ under standing conditions, they can be observed when cells are exposed to shear forces via shaking of the culture. Furthermore, *P. aeruginosa* Δ*phz* mutants are inhibited in their ability to form biofilms under anaerobic, shaking conditions. For *H. volcanii*, we could show for the first time that biofilms can form under shaking conditions and exhibit a different biofilm architecture than under standing conditions. Additionally, *H. volcanii* archaella seem to be involved in the initial surface adhesion under shaking, but not under standing, conditions. These results show not only the biological insights that can be gained from biofilm assays under a variety of conditions but also highlight the versatility of the mPAD mount. With the increased availability of 3D-printing capabilities and the straightforward changes to the design of the mPAD, the range of applications will widen even further.

## Acknowledgements

We would like to thank the University of Pennsylvania Libraries’ Biotech Commons for courtesy 3D-printing of mPAD mounts. S.S., H.S., and M.P acknowledge support from the National Science Foundation Grant 1817518. K.C. acknowledges support from a Young Investigator Award from the Army Research Office, grant number W911NF-19-1-0024.

## Conflict of interest

The authors declare no conflict of interest.

## Supplemental Material

Supplemental Figure 1: 3D-printing of the mPAD mount allows for straightforward changes to its design.

